# Target-Specific Discovery of BMM_1567 Restores Aminoglycoside Activity Against Multidrug-Resistant Gram-Negative ESKAPE Pathogens

**DOI:** 10.64898/2026.07.20.739709

**Authors:** Meenal Chawla, Lekshmi Narendrakumar, Deepjyoti Paul, Ramani Shyam Kapuganti, Shantanu Kumar, Debaleena Das, Kajal Kamboj, Susmita Bakshi, Payal Priyadarshi, Dinesh Mahajan, Shailendra Asthana, Bhabatosh Das

**Affiliations:** Microbial Research Centre, BRIC-Translational Health Science and Technology Institute, NCR Biotech Science Cluster, Faridabad – 121001, India; Computational and Mathematical Biology Centre, BRIC-Translational Health Science and Technology Institute, NCR Biotech Science Cluster, Faridabad – 121001, India; Drug Discovery Center, BRIC-Translational Health Science and Technology Institute, NCR Biotech Science Cluster, Faridabad – 121001, India

## Abstract

The global emergence of multidrug-resistant (MDR) ESKAPE pathogens has significantly reduced the effectiveness of existing antibiotics, highlighting the urgent need for new strategies to restore antimicrobial susceptibility. Here, we report the discovery and mechanism of BMM_1567, a peptide potentiator that enhances aminoglycoside efficacy against MDR pathogens. A genetically defined reporter-based screen identified BMM_1567 as a potent inhibitor of aminoglycoside resistance, potentiating spectinomycin activity against MDR Gram-negative ESKAPE isolates at low micromolar concentrations. Structural modeling and molecular dynamics simulations indicated that BMM_1567 interacts with residues lining the antibiotic-binding groove of aminoglycoside-modifying enzymes (ANT, APH, AAC), with highest affinity for ANT (ΔG_bind = –62.25 kcal/mol), suggesting competitive inhibition of substrate binding. Site-directed mutagenesis of key ANT residues identified critical amino acids involved in BMM_1567 binding, confirming their role in mediating spectinomycin potentiation. In murine abscess model using XDR *E. coli*, BMM_1567 in combination with spectinomycin significantly reduced bacterial burden and pro-inflammatory cytokine levels, comparable to colistin. Collectively, these findings establish BMM_1567 as a promising aminoglycoside potentiator that restores antibiotic activity against MDR pathogens through direct inhibition of resistance enzymes, while exhibiting in vivo efficacy and a remarkably low propensity for resistance development.

## Introduction

The global surge in antimicrobial resistance (AMR) has emerged as a critical threat to modern medicine, particularly due to the increasing prevalence of multidrug-resistant (MDR) Gram-negative bacteria within the ESKAPE group *Enterococcus faecium, Staphylococcus aureus, Klebsiella pneumoniae, Acinetobacter baumannii, Pseudomonas aeruginosa*, and *Enterobacter* species (1–3). These pathogens are notorious for their capacity to evade existing antibiotic therapies, leading to prolonged infections, higher treatment costs, and increased mortality (4–8). The inadequacy of current antibiotics to combat these resilient pathogens has prompted urgent calls for novel therapeutic strategies that either restore or enhance the efficacy of existing drugs (9–11).

One promising approach to combat AMR is the use of antibiotic potentiators, agents that possess little or no intrinsic antibacterial activity but can restore or enhance the efficacy of conventional antibiotics against resistant bacterial pathogens (12, 13). These agents function by targeting and impairing bacterial resistance mechanisms, thereby restoring antibiotic efficacy (14). Clinical success with β-lactamase inhibitors has established a proof-of-concept for such approaches (15, 16). However, despite the clear therapeutic value, only a limited number of antibiotic-potentiator combinations have been translated into clinical practice (13). This underscores the need for comprehensive, mechanism-guided discovery pipelines capable of identifying and validating effective potentiators, particularly for antibiotic classes such as aminoglycosides, which are often rendered ineffective due to acquired enzymatic resistance (17, 18).

In this study, we aimed to discover novel potentiators that enhance the efficacy of aminoglycosides against MDR bacteria, including clinically derived ESKAPE isolates. Employing a high-throughput screening (HTS) platform and a phenotypic, cell-based assay using genetically engineered reporter strains harbouring resistance determinants from diverse bacterial taxa, we screened a library of 4,417 commercially available compounds. This approach allowed for the rapid and targeted identification of compounds that could potentiate aminoglycosides activity by suppressing the activity of aminoglycoside modifying enzymes. From the initial HTS, several candidate compounds were identified as potentiators. Of these, BMM_1567 emerged as a lead compound due to its consistent ability to restore the activity of aminoglycoside antibiotics against multiple aminoglycoside-resistant reporter strains, including *E. coli* strains expressing the *aadA1*, *aph3’*, and *aac3’* resistance genes. This effect extended beyond laboratory strains, showing broad-spectrum potentiation activity against clinical MDR isolates of *E. coli*, *A. baumannii*, *K. pneumoniae*, and *P. aeruginosa*. The therapeutic potential of BMM_1567 was further validated in vivo using a murine skin abscess model infected with an XDR *E. coli* clinical isolate, where bacterial clearance levels with BMM_1567 and aminoglycoside antibiotic is comparable to colistin, a last-resort antibiotic. To further validate the mechanistic basis of action, we performed in silico analysis for uncovering potential interactions site with the target molecule, site-directed mutagenesis (SDM) of key amino acid residues within the ANT enzyme.

Taken together, our findings highlight BMM_1567 as a promising aminoglycoside potentiator with both broad-spectrum and clinically relevant activity. Its low effective concentration, and ability to restore activity against MDR ESKAPE pathogens, make it a strong candidate for further preclinical development. Moreover, our integrated discovery platform combining phenotypic screening using specific and sensitive reporter strains, computational modelling, molecular validation, and in vivo efficacy testing offers a robust pipeline for the identification of future antibiotic adjuvants aimed at overcoming the ever-growing threat of antibiotic resistance.

## Results

### Development of Chromosomally Integrated Reporter Strains for High-Throughput Discovery of Aminoglycoside Potentiators

To enable the systematic discovery of aminoglycoside potentiators targeting clinically relevant resistance determinants, we established a panel of genetically defined reporter strains optimized for high-throughput phenotypic screening. An ideal screening platform should combine high sensitivity with target specificity while ensuring efficient intracellular access of test compounds. To achieve this, we selected *V. cholerae* N16961 and *E. coli* ATCC 25922 as reporter backgrounds . *V. cholerae* N16961 is intrinsically susceptible to a broad range of antibiotics, including aminoglycosides, and possesses a relatively permeable outer membrane that facilitates enhanced uptake of small molecules (19, 20). This characteristic increases the likelihood of detecting weakly active compounds that may otherwise remain undetected in conventional Gram-negative screening systems. To prioritize resistance determinants for reporter strain construction, we performed a comparative genomic analysis of Indian and global clinical isolates of Gram-negative ESKAPE pathogens using whole-genome sequences (n=66,039) retrieved from the NCBI database (**Fig. 1A**). This analysis identified nine highly prevalent aminoglycoside resistance genes representing the three major classes of aminoglycoside-modifying enzymes (AMEs): aminoglycoside phosphotransferases (APHs), aminoglycoside acetyltransferases (AACs), and aminoglycoside nucleotidyltransferases (ANTs) (**Fig. 1A**). These enzymes collectively account for a substantial proportion of aminoglycoside resistance observed in clinically important Gram-negative pathogens. To create a genetically stable and mechanistically defined screening platform, each resistance determinant was irreversibly integrated into the chromosome of *V. cholerae* N16961 or *E. coli* ATCC 25922 (21, 22). Chromosomal insertion was achieved at the conserved *dif* locus using our previously developed XerC-XerD-mediated site-specific recombination system, ensuring stable inheritance and minimizing variability associated with plasmid-based expression (23). To maintain physiologically relevant and reproducible expression levels, all resistance genes were placed under the control of the constitutive *htpG* promoter (24).

**Figure 1:**
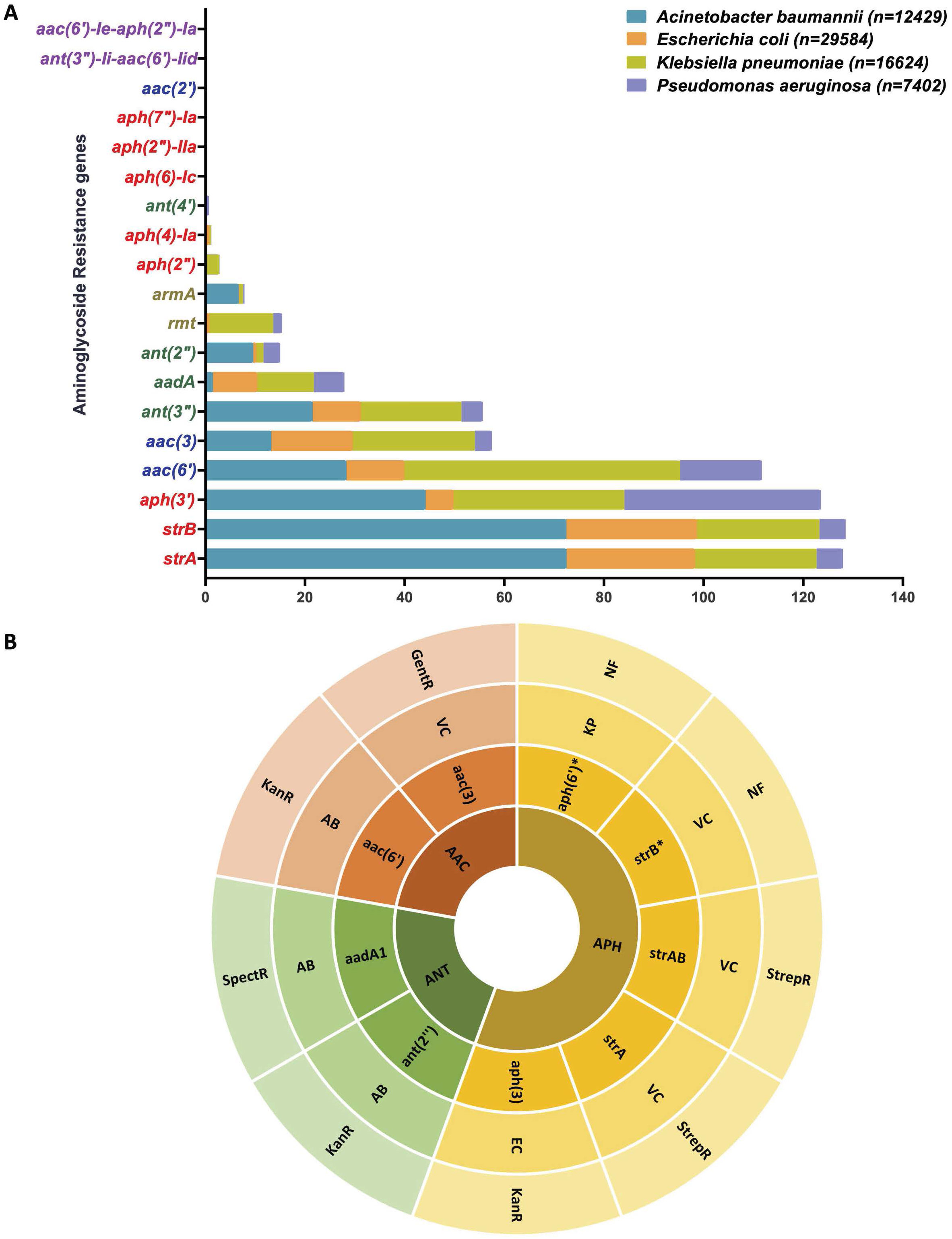
Prevalence of aminoglycoside resistance genes and development of aminoglycoside reporter strains. **A)** The prevalence of aminoglycoside resistance genes in Gram-negative members of ESKAPE pathogens. **B)** Sunburst diagram of distinct aminoglycoside resistance genes in reporter strains. The inner segment indicates distinct aminoglycoside subclasses, the second segment represents specific resistance genes associated with each subclass, third ring represents the clinical isolate from which gene was amplified and outer segment shows the functionality for aminoglycoside antibiotics. (VC – *V. cholerae*, EC - *E. coli*, AB **-** *A. baumannii,* KP *- K. pneumoniae,* Strep - Streptomycin, Kan – Kanamycin, Spec – Spectinomycin, NF – Non-functional).

Using this strategy, we generated a library of sixteen engineered reporter strains harboring aminoglycoside resistance genes derived from diverse clinical isolates (**Table 1**). Antimicrobial susceptibility profiling confirmed that each engineered strain exhibited the expected resistance phenotype toward its cognate aminoglycoside substrate, validating both the functionality of the integrated resistance determinants and the robustness of the reporter platform (**Fig. 1B**; **Tables 1 and 2**). Collectively, these strains constitute a comprehensive, genetically stable, and target-specific screening resource for the identification of compounds capable of disabling aminoglycoside resistance mechanisms and restoring antibiotic activity against multidrug-resistant Gram-negative pathogens.

**Table 1:**
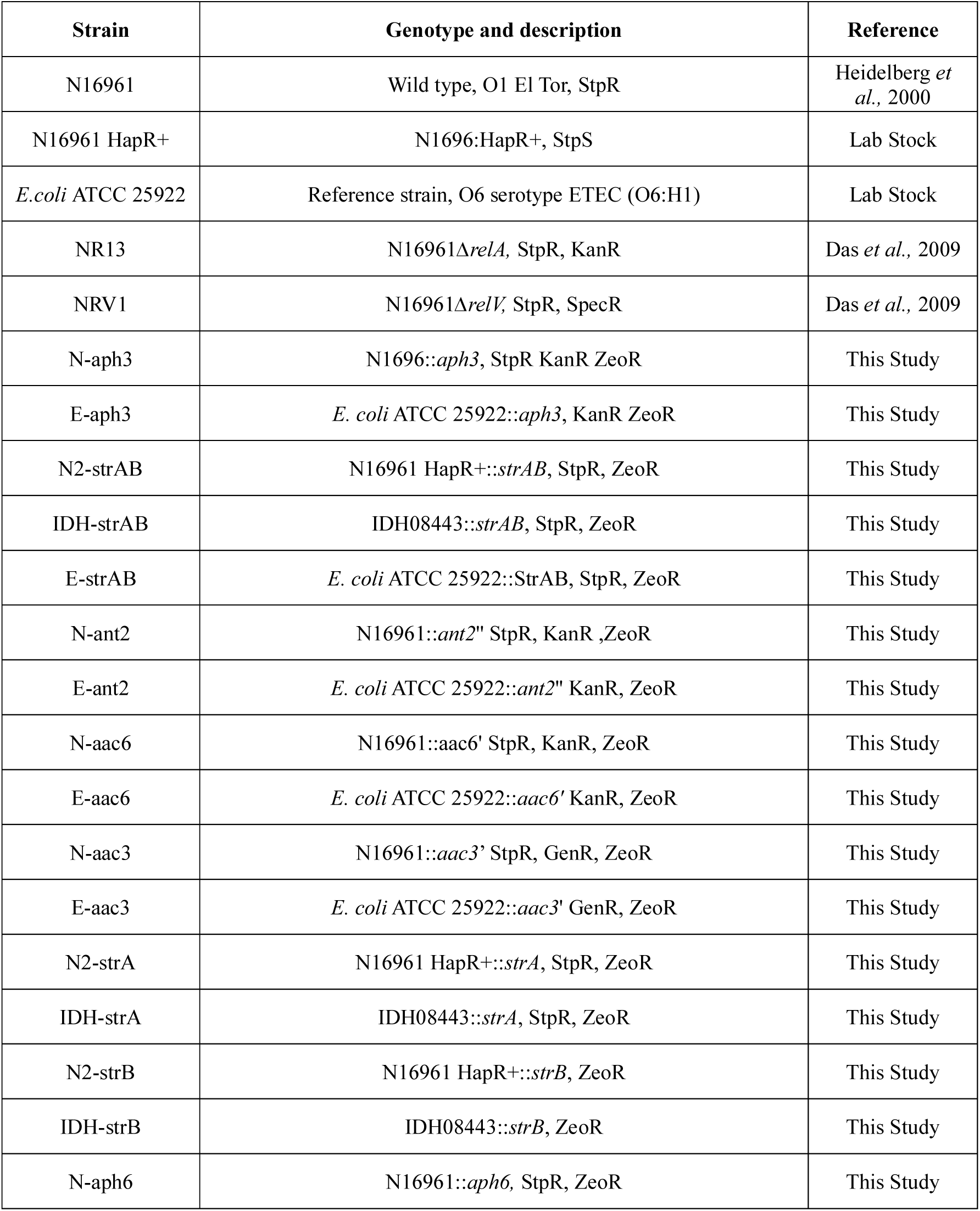
Genotype and Phenotype Characteristics of Wild-Type and Genetically Modified Strains Used in the Study.

**Table 2:**
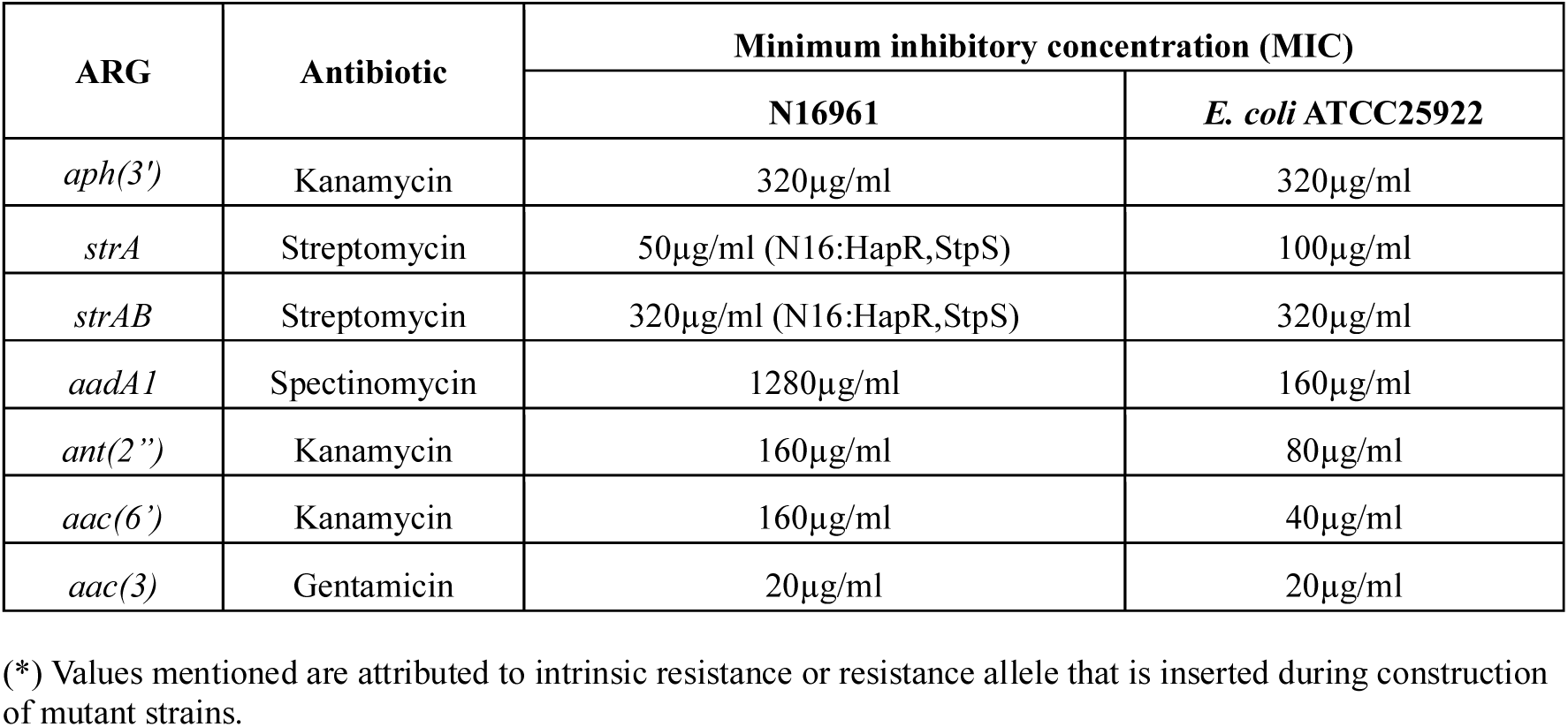
Minimum Inhibitory Concentration (MIC) of Various ARGs in Reporter Strains N16961, and *E. coli* ATCC 25922.

### Identification of Aminoglycoside Potentiators Against Resistant Bacteria

Aminoglycoside potentiator screening was done using two commercially available libraries: the Selleckchem Natural Compound Library (n = 803) and the MedChem Express (MCE) Library (n = 3614). Most compounds in the MCE library are either FDA-approved or currently undergoing various phases of clinical trials. Primary screening was performed at 10 µM concentration for all the compounds (n = 4417). The compound libraries were screened using *V. cholerae* N16961 derived reporter strain, N-aph3 which is resistant to kanamycin antibiotic. In the primary screening, 326 compounds exhibited greater than 30% growth inhibition against the Naph3 reporter strain and were therefore selected as preliminary hits for further evaluation **(Fig. 2)**. Among the 326 active compounds recognized, 201 were well-known antibiotics, 91 include compounds with reported antibacterial activity in the literature, and 34 represented novel compounds with no prior reports of antibacterial activity **(Fig. 3A)**. Among these 34 compounds, eight compounds displayed synergistic or potentiating effects ranging from 35 to 90% growth inhibition with kanamycin in the Naph3 reporter strain **(Fig. 3B and 3C)**. Eight potentiating or synergistic compounds were further evaluated in *E. coli* reporter strain Eaph3.

**Figure 2:**
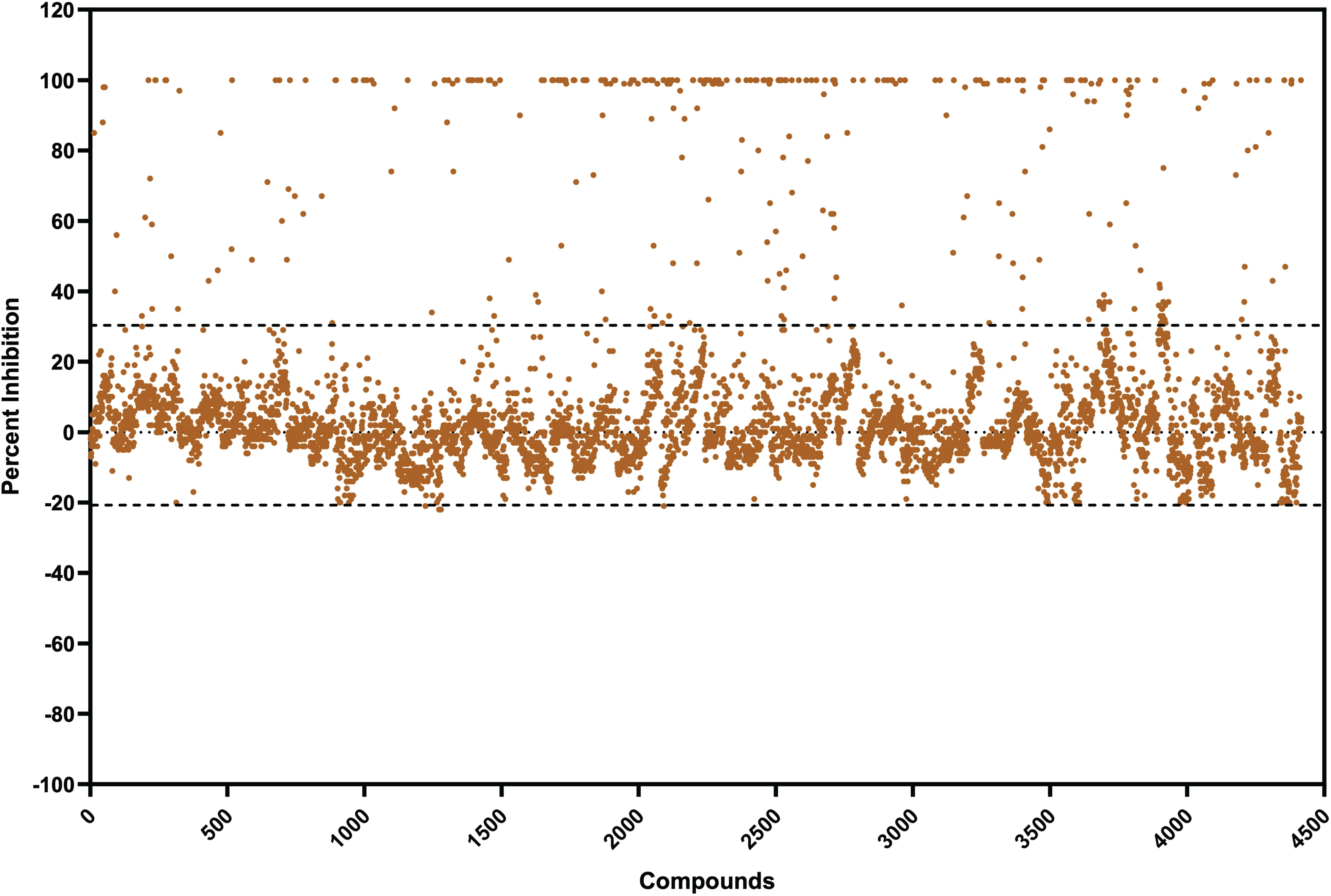
Primary screening of compounds against reporter strains. A scatter plot representing primary screening of 4417 compounds, comprising 3614 compounds from FDA-approved (MedChem) compound library and 803 compounds from Selleckchem natural compound library. The screening was conducted against *aph(3’)* using Naph3 reporter strains. The antibiotics employed for screening were kanamycin.

**Figure 3:**
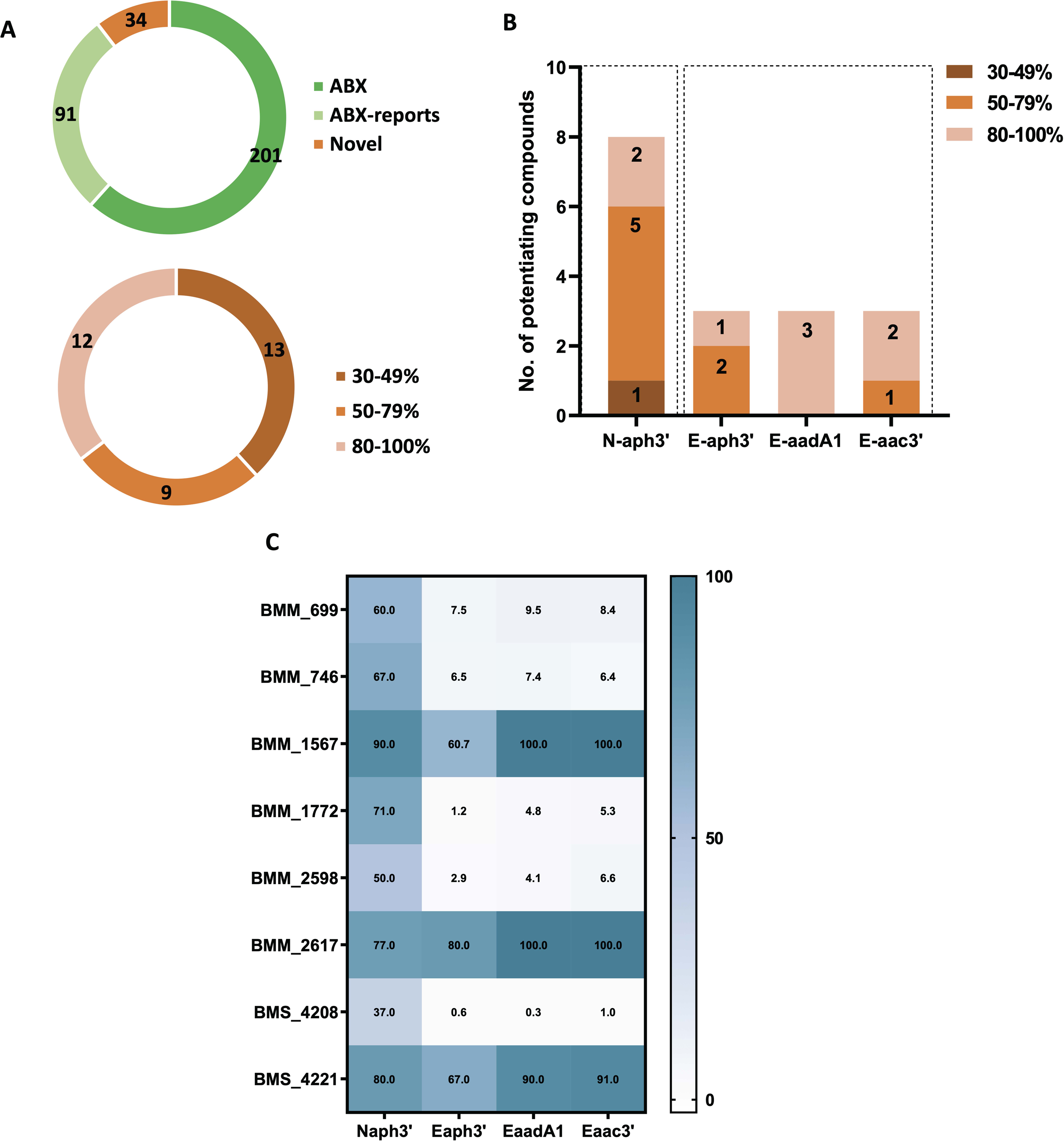
Secondary screening of compounds with and without antibiotics. **A)** Donut chart illustrating the classification of active compounds from primary screening into three categories: known antibiotics, literature reported antibiotic-like compounds, and novel compounds. Further the distribution of 34 novel compounds based on their percent inhibition of bacterial growth. **B)** Number of active compounds against specific alleles in N16961 and *E. coli* ATCC reporter strains. **C)** Heatmap representing the bacterial growth inhibition in presence of combination of compounds with antibiotics against Naph3’, Eaph3’, EaadA1, and Eaac3’ reporter strain. Kanamycin was used for Naph3’ and Eaph3’ strains, spectinomycin for EaadA1 and gentamicin for Eaac3’ reporter strains. The color scale indicates the percent inhibition values. Percent inhibition data represent biological triplicates as mean.

Only three of these compounds BMM_1567, BMS_4221 and BMM_2617 demonstrated activity in Eaph3 **(Fig. 3C)**. These three compounds were also assessed in EaadA1 and Eaac3 reporter strains with spectinomycin and gentamicin antibiotics, respectively **(Fig. 3C)**. To eliminate the possibility of library-specific artifacts, the identified hit compounds were repurchased from independent commercial vendors and were observed to reproduce their initial activity. The best combinations of compound and antibiotic were determined in *E. coli* reporter strain through checkerboard assay **(Supplementary Fig. 1)**. These three compounds displayed activity with the three tested aminoglycoside antibiotics but at different concentrations. The compound BMS_4221 displayed potentiating activity at concentrations ranging from 50-200µM. The compound BMM_1567 exhibited activity at concentrations ranging from 2.5-20µM, showing potentiating effect at lower concentrations and synergistic effect at higher concentrations. The third compound BMM_2617, a β-lactam antibiotic, displayed a potentiating effect at concentration of 0.312µM **(Supplementary Fig. 1)**. The structures of aminoglycoside potentiators identified in Naph3 reporter strains is shown in **Supplementary Fig. S2**.

### Evaluation of aminoglycoside potentiating compounds against MDR clinical isolates

Aminoglycoside potentiating compounds showed promising activity, particularly with spectinomycin, in enhancing its efficacy against resistant bacteria, including EaadA1 strain. To further evaluate their clinical potential, we tested these compounds in combination with spectinomycin against Gram-negative members of the ESKAPE pathogens. The selected MDR clinical isolates, exhibit resistance to over 10 different antibiotics and contain various antibiotic resistance cassettes (**Supplementary Table S1)**. In our assay, BMM_1567 exhibited a strong potentiating effect with spectinomycin, achieving 50-100% potentiation at low concentrations (1.25 µM to 5 µM) against *E. coli*, *A. baumannii, K. pneumoniae*, and *P. aeruginosa* MDR clinical isolates **(Fig. 4A-D, Supplementary Table S1)**. Similarly, BMS_4221 showed inhibition ranging from 50-100% at higher concentrations (50µM to 200 µM) against *E. coli* and *K. pneumoniae* isolates, minimal activity against *P. aeruginosa* but no activity was observed against *A. baumannii* isolates **(Supplementary Table S1, Supplementary Fig. 3A-D)**. On the other hand, BMM_2617 did not show substantial inhibition against any of the tested isolates, likely due to the presence of multiple β-lactam resistance genes in these clinical isolates. The activity of BMM_1567 was further validated by colony-forming unit (CFU) assays, confirming its potential as the most promising potentiating compound. We extended its evaluation with additional aminoglycosides, such as gentamicin and kanamycin, against the same MDR clinical isolates. When combined with gentamicin, BMM_1567 did not exhibit significant activity against *K. pneumoniae* isolates. However, it displayed potentiating or synergistic activity (30-100% inhibition) against *E. coli* and *A. baumannii* at a concentration of 2.5 µM **(Supplementary Table S1, Supplementary Fig. S3E and S3F).** When tested with kanamycin, BMM_1567 demonstrated a potent inhibitory effect (95%) against MDR *K. pneumoniae* isolates at the same concentration (2.5 µM) **(Supplementary Table S1, Supplementary Fig. S3G)**, and 50-90% inhibition was observed against *A. baumannii* isolates, though at a higher concentration of 20 µM **(Supplementary Table S1, Supplementary Fig. S3H)**. The BMM_1567 with kanamycin has also shown potentiation against MDR *Enterobacter roggenkampii* isolate (**Supplementary Table S1**).

**Figure 4:**
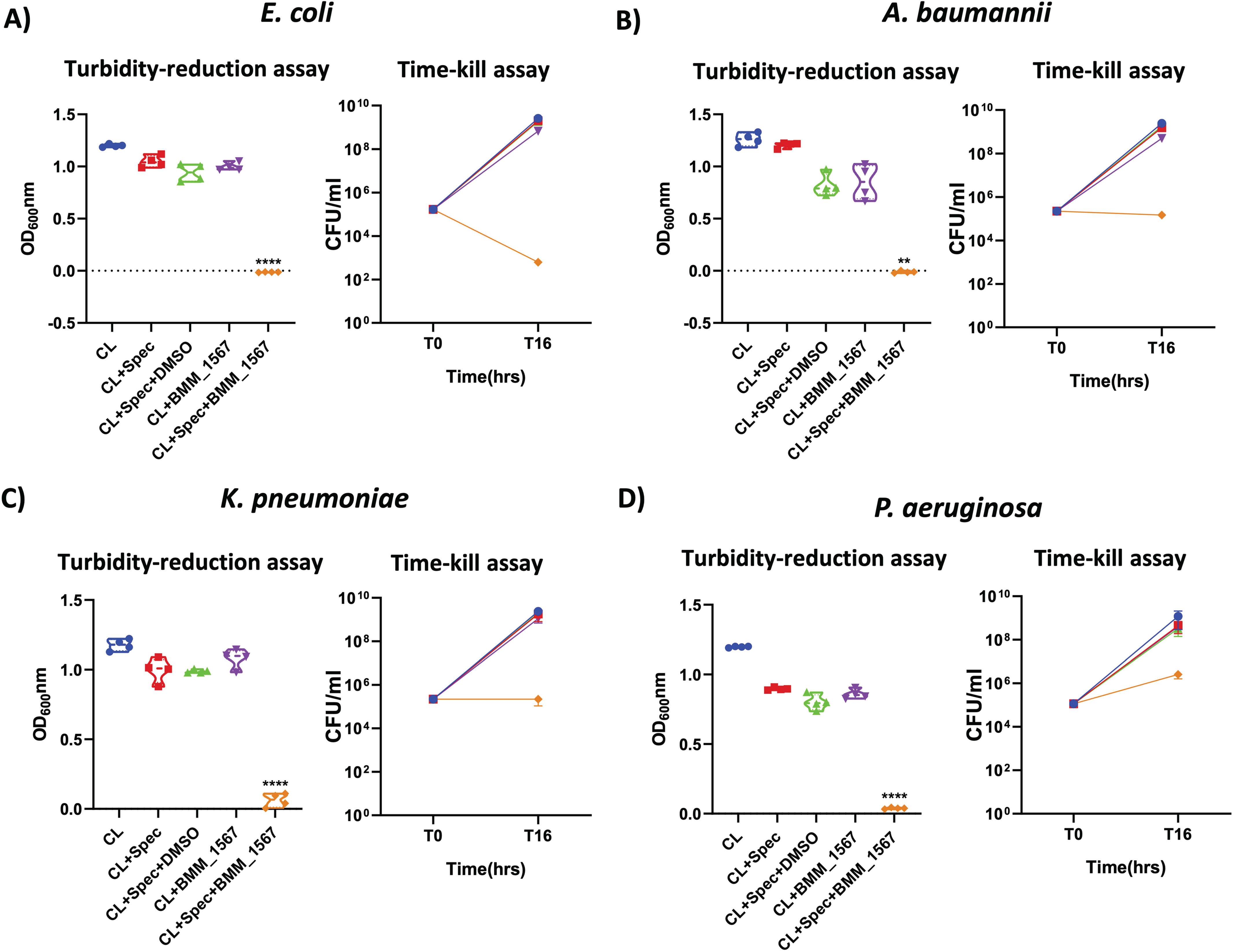
Evaluation of Active Potentiating Compounds Against Gram-Negative ESKAPE Pathogens. Assessed the activity of BMM_1567 with spectinomycin antibiotics against: **A)** *E. coli*, **B)** *A. baumannii,* **C)** *K. pneumoniae,* and **D)** *P. aeruginosa*. Bacterial killing was quantified using 16h turbidity reduction assays and time kill assays with plating on LA agar plates. Antibiotic concentrations used are 40µg/ml spectinomycin. For BMM_1567, concentrations used were 5µM for *K. pneumoniae* (FGL-174) and *P. aeruginosa* (FGL-31), and 2.5µM for *A. baumannii* (FGL-166) and *E. coli* (FGL-155). Turbidity reduction and time kill assay data are biological triplicates shown as mean ± SD. Unpaired t-test with Welch’s correction was used to determine statistical significance (P < 0.05), with significance calculated relative to the compound alone (p-values <0.05 (*), <0.01 (**), p-values <0.001 (***) and p-values < 0.0001 (****). Strains utilized were MDR clinical isolates harboring ARGs. (CL - Bacterial culture, Spec – Spectinomycin).

### Validation of a Lead Antibiotic Potentiator in a Murine Infection Model

To assess the in vivo potentiating activity of the hit compounds, we used a murine skin abscess model infected with a clinically derived XDR *E. coli* isolate recovered from a pus sample. A localized skin infection with XDR *E. coli* was established (**Fig. 5A**). The potentiating effect of BMM_1567 in combination with spectinomycin and effect of BMS_4221 with spectinomycin was assessed, allowing us to determine the therapeutic potential of antibiotic-potentiator pairing in a relevant pre-clinical context. Single dose in every 24h was delivered to the infected area. Two days post-infection (two doses), treatment with BMM_1567 with spectinomycin resulted in a significant reduction in bacterial load, by 2 to 3 orders of magnitude - compared to the untreated control, spectinomycin alone, vehicle control, and BMM_1567 monotherapy groups. Its potency was comparable to the activity observed in mice group treated with colistin (**Fig. 5B**). However, the combination of BMS_4221 with spectinomycin did not show significant effect. Essentially, no significant changes in body weight were observed with any of the combination evaluated in the in vivo study (**Fig. 5C**). The bacterial isolates from the BMM_1567 with spectinomycin group were re-evaluated in vitro to assess the potentiating activity of BMM_1567 in combination with spectinomycin. The results were compared with the original bacterial isolate, and no change in potentiating activity was observed **(Supplementary Fig. S4A)**. Additionally, we assessed systemic inflammation by measuring IL-6 and IL-1β expression levels in blood across treatment groups. Compared to the untreated group, both the BMM_1567 with spectinomycin combination and colistin treatment significantly reduced the expression of these pro-inflammatory cytokines (**Supplementary Fig. S4B and S4C**). These findings are promising, highlighting that the potentiator-antibiotic combination exerts a substantial therapeutic effect at low concentrations, even after infection is established.

**Figure 5:**
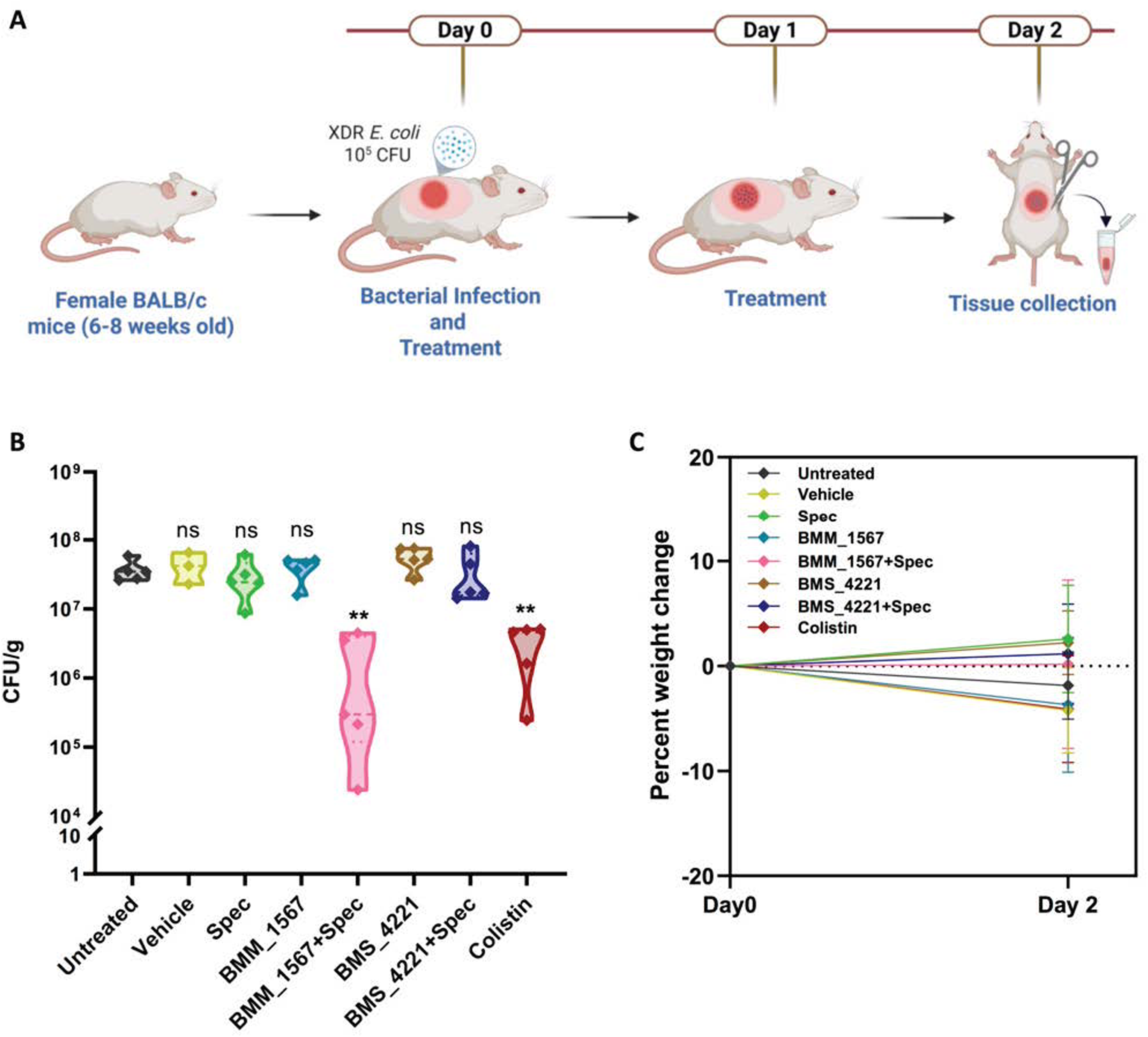
Assessment of potentiating activity of lead compound with antibiotic in preclinical mouse model. **A)** Schematic of mouse skin abscess model for in vivo activity assessment. Mice incision was inoculated with XDR *E. coli* strain. Treatments were administered 1h post-infection (day0) and again at day 1. Skin samples were collected at day 2 for bacterial counts. **B)** Bacterial count of the different experimental groups of the mice after 2 days post infection. Five mice per group, except for the vehicle group which included three mice. The data were log_10_ transformed and statistical significance was determined using one-way ANOVA followed by Dunnett’s multiple comparison test. All the groups were compared to the untreated group. **C)** Body weight of mice from different experimental groups measured on Day 0 and Day 2. No significant changes were observed across the groups during the experimental period. Data are presented as mean ± standard deviation (SD).

### Elucidating the Molecular Mechanism of Potentiator-Mediated Inhibition

BMM_1567 demonstrated the most promising potentiating activity in combination with aminoglycoside antibiotics, both in vitro and in a murine skin abscess model. Therefore, its mechanism of action was further investigated. BMM_1567 alone exhibited no intrinsic antimicrobial activity; however, when combined with aminoglycosides, it produced significant growth inhibition. These findings suggest that BMM_1567 likely inhibits the activity of the resistance enzyme, thereby restoring the antibacterial efficacy of aminoglycoside antibiotics. To investigate this hypothesis further, we performed the in-silico analysis to assess the binding of BMM_1567 to the ANT, APH, and AAC enzyme encoded by *aadA1*, *aph3’* and *aac3’* gene, respectively. SiteMap analysis coupled with unbiased molecular docking identified the antibiotic-binding groove as the most energetically favorable binding pocket for BMM_1567 in all three aminoglycoside-modifying enzymes, suggesting a shared mechanism of inhibition (**Supplementary Fig. S5A**). The subsequent MD simulations were done to measure the structural stability and dynamical changes of each complex. The root means square deviation (RMSD) of ANT revealed that both the APO (ANT) and complex (ANT + BMM_1567) achieved stability after the 50ns of trajectory and with respect to APO, the ANT-BMM_1567 complex and ANT-spectinomycin complex seems stable. The RMSF analysis presented similar fluctuation patterns across all three systems **(Fig. 6A)**. Further interaction fingerprinting of most stable complex revealed some key residues in ANT, D43, E83, W108, and N181, which were common with spectinomycin and BMM_1567, confirming overlapped binding site of the two (**Fig. 6B and Supplementary Fig. S5B**) (**Table 3**). The average ΔG_bind_ of BMM_1567 with ANT was -62.25 kcal/mol while ΔG_bind_ for ANT-spectinomycin was -34.15 kcal/mol (**Fig. 6C**).

**Figure 6:**
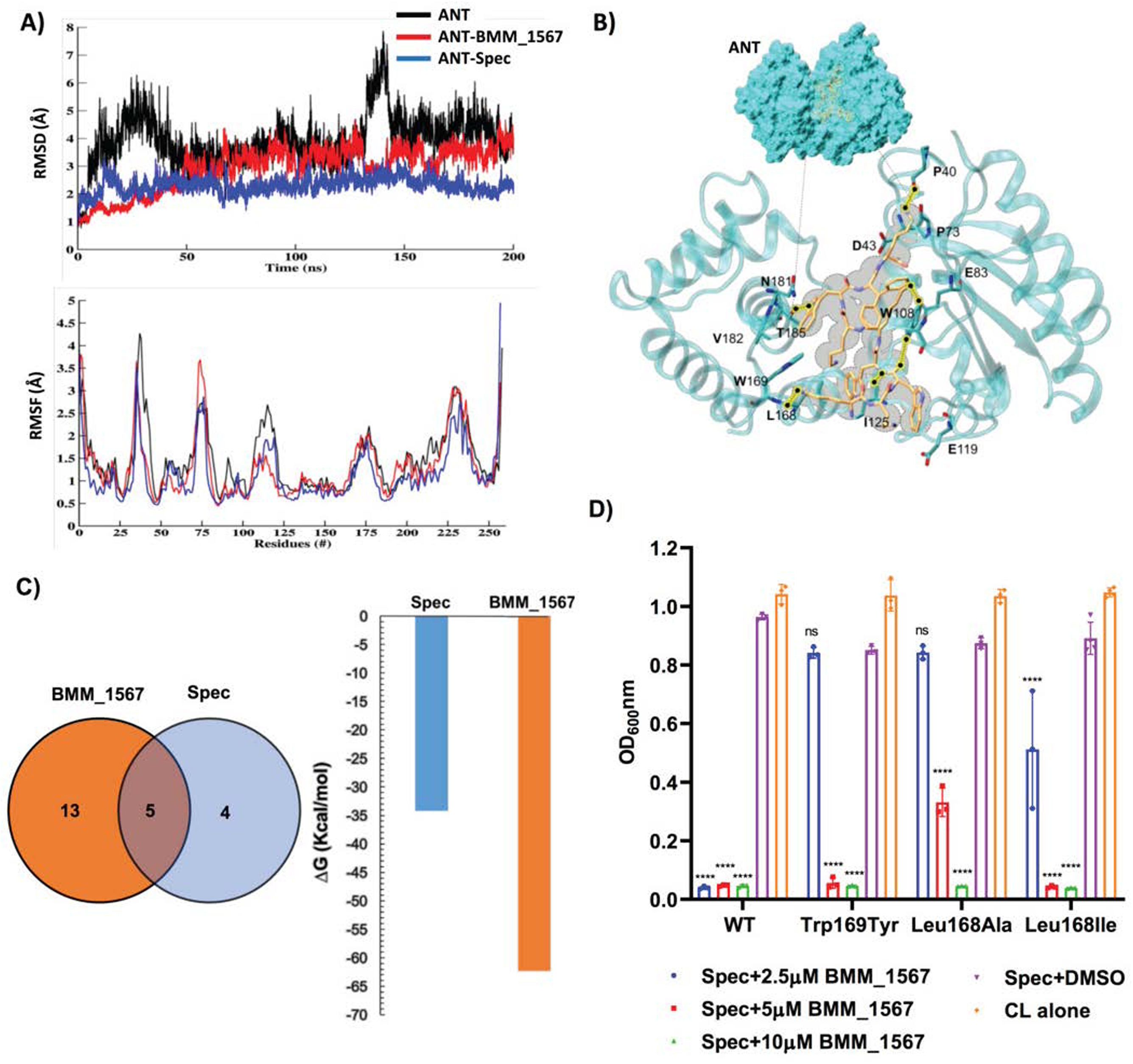
Mechanistic insights into BMM_1567 binding with ANT enzyme: molecular docking, dynamic simulations, and SDM validation. **A)** Time-dependent dynamical changes monitoring through molecular dynamics simulations - the RMSD and RMSF of the ANT systems (Apo, ANT-BMM_1567 complex and ANT-Spec Complex). **B)** The most stable states of BMM_1567 in the binding site of ANT and the interaction map between binding site residues and BMM_1567. The hydrogen-bonds are shown by dotted line and black in color. **C)** Venn diagram of comparative analysis of interacting residues in ANT-BMM_1567 and ANT-Spec complex and the quantitative binding energy between the complexes, shown in bar graph. **D)** Effect of SDM mutations on potentiating activity of BMM_1567. Wild-type (WT) and mutant SDM strains were grown in the presence of spectinomycin and varying concentrations of the BMM_1567. The Growth assessment (OD₆₀₀) was measured after 16h of incubation. Data represent the mean ± SD of three biological replicates. Statistical significance was determined using two-way ANOVA followed by Dunnett’s multiple comparisons test, comparing each group with Spec+DMSO control.

**Table 3:**
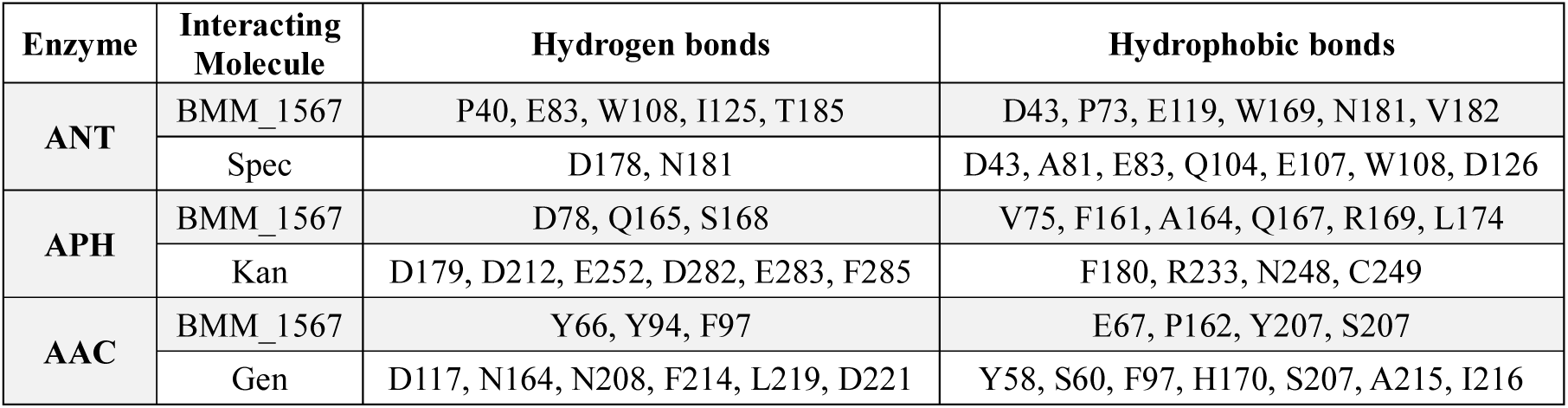
Hydrogen bonding and hydrophobic interactions between different enzymes and interacting molecules.

The MD simulations of APH and AAC indicated that APH-BMM_1567 complex is the most stable system, while AAC shows similar dynamics in APO, BMM_1567-bound, and gentamicin-bound forms (**Fig. 7A and 7D)**. The key interacting residues with favorable energetic contributions were mapped for APH and AAC as well **(Table 3)**. The BMM_1567 showed stronger binding than the antibiotics, with average ΔG_bind_ of −58.35 kcal/mol vs −35.35 kcal/mol for kanamycin with APH, and −55.44 kcal/mol vs −37.28 kcal/mol for gentamicin with AAC **(Fig. 7B-C, and Fig. 7E-F**). Overall, the computational analysis indicates that BMM_1567 is stable in APH, ANT and AAC, however, the best binding affinity was observed in ANT, followed by APH and least in AAC.

**Figure 7:**
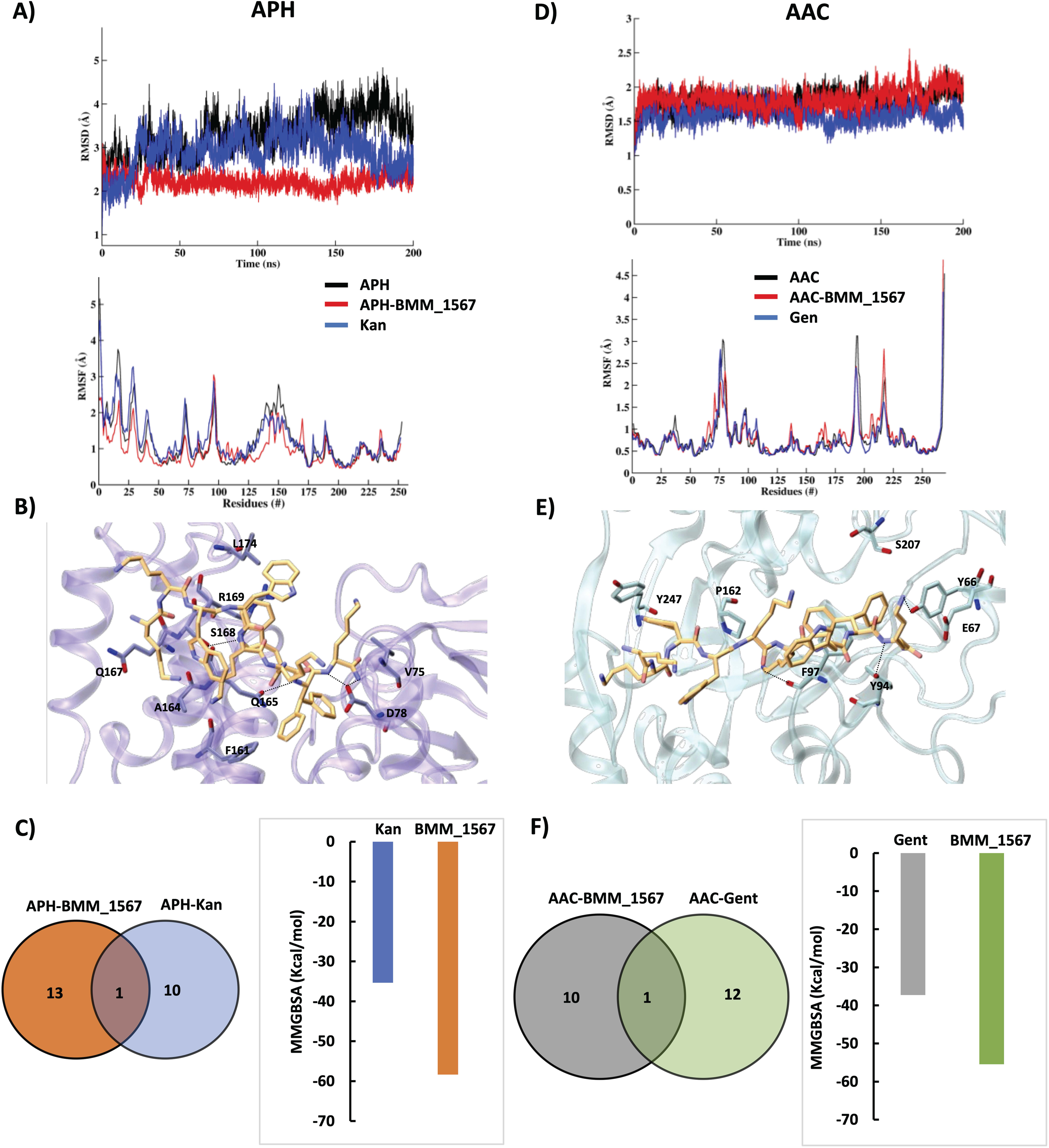
BMM_1567 binding with APH and AAC enzyme: Molecular docking and dynamic simulations. **A)** The RMSD and RMSF of the APH systems (Apo(APH), APH-BMM_1567 Complex and APH-Kan Complex). **B)** The interaction map of APH-BMM_1567 complex. The residues of APH binding site with most favorable energetic contribution and interacting peptide is shown in licorice and color atom-wise (C:violet/orange, O:red, and N:blue). H-bond are shown is black dotted line. The APH is rendered in cartoon transparent view. **C)** Venn diagram of comparative analysis of interacting residues in APH-BMM_1567 and APH-Kan complex. The quantitative binding energy between the APH-BMM_1567 and APH-Kan complexes, shown in bar graph. **D)** The RMSD and RMSF of the AAC systems (Apo(AAC), AAC-BMM_1567 complex and AAC-Gen Complex). **E)** The interaction map of AAC-peptide complex. The residues of AAC binding site with most favorable energetic contribution and interacting peptide is shown in licorice and color atom-wise (C:sky blue/orange, O:red, and N:blue). H-bond are shown is black dotted line. The AAC is rendered in cartoon transparent view. **F)** Venn diagram of comparative analysis of interacting residues in AAC-BMM_1567 and AAC-Gen complex. The quantitative binding energy between the AAC-BMM_1567 and AAC-Gen complexes, shown in bar graph. (Kan-Kanamycin, Gen-Gentamicin).

Following the in-silico confirmation of BMM_1567 binding with aminoglycoside resistance enzyme, the in vitro validation was conducted to confirm the structure dependent action of BMM_1567. Site-directed mutagenesis (SDM) was employed for the same by targeting ANT enzyme, due to its superior potentiation effect observed in vitro and stronger binding affinity observed in silico. Notably, the ANT residues interacting with BMM_1567 were either identical to or located adjacent to those involved in spectinomycin binding. To preserve the resistance activity of the ANT enzyme while investigating ANT-BMM_1567 interaction, SDM was strategically performed on residues unique to BMM_1567 binding. From per-residue analysis, residue Leu168 and Trp169 was identified as major contributing residues and therefore they were selected for mutational studies via computational alanine scanning (**Supplementary Fig. S5B)**. The Leu168Ala (Alanine) and Trp169Ala substitutions resulted in reduced resistance activity of the ANT enzyme, compared to the wild-type (WT) enzyme. The observed loss of resistance is likely due to their location within the spectinomycin binding pocket, inadvertently affecting resistance functionality of ANT enzyme. Further to retain structural and chemical similarity, in-silico analyses suggest Trp169Tyr (Tyrosine) and Leu168Ile (Isoleucine) mutations as suitable, which were subsequently done. Trp169Tyr mutation has shown better resistance activity as compared to Trp169Ala mutation. However, the level of resistance in Leu169Ala and Leu168Ile mutations were similar. The effect on the potentiating activity of BMM_1567 with spectinomycin was determined for all the ANT mutations (**Fig. 6D**). To take into account the effect of different resistance activity, the FICI values were calculated. Compared to WT enzyme (FICI = 0.1875), Leu168Ala has shown much higher FICI (0.375) and Leu168Ile and Trp169Tyr showed FICI values of 0.25, suggesting Leu168Ala substitutions has much larger effect on potentiating activity of BMM_1567 **(Table 4)**. We further assessed the frequency of resistance development to the combination of BMM_1567 and spectinomycin. However, when the EaadA1 strain was plated on medium containing 4× MIC of BMM_1567 in combination with spectinomycin, no true resistant colonies were recovered. We therefore estimate resistance frequency to be <1 × 10⁻^8^.

**Table 4:**
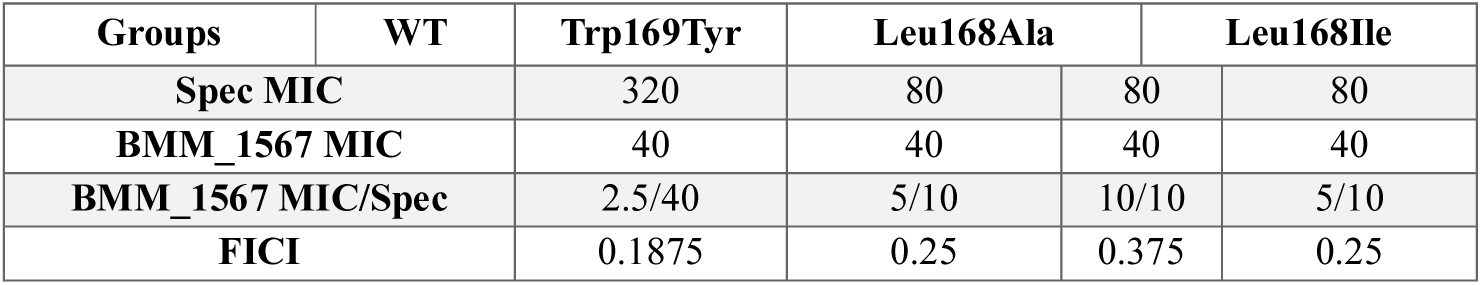
Fractional Inhibitory Concentration Index (FICI) values for wild-type (WT) and SDM mutant strains of ANT enzyme tested with the spectinomycin and BMM_1567.

## Discussion

The emergence and proliferation of MDR Gram-negative pathogens represent a pressing global health crisis (3,25). In particular, the widespread dissemination of aminoglycoside resistance mechanisms, often mediated by enzymatic modification of antibiotics, has significantly reduced the clinical utility of this important antibiotic class (26–28). The global rise in resistance to β-lactams and β-lactam/β lactam-potentiator combinations has renewed interest in aminoglycosides as an effective therapeutic agent against MDR pathogens (17). However, the growing prevalence of aminoglycoside resistance and the absence of clinically potent aminoglycoside potentiators have severely hindered their emergence as a frontline strategy in combating MDR infections (29–31). Our findings demonstrate that inhibition of aminoglycoside-modifying enzymes represents an effective strategy to restore aminoglycoside susceptibility in multidrug-resistant bacteria, leading to the identification of novel antibiotic adjuvants with potentiating activity. By combining target-guided phenotypic screening using genetically defined aminoglycoside-resistance reporter strains with synergy testing, in vitro and in vivo validation, and computational modeling, we identified multiple aminoglycoside potentiators, including the lead molecule BMM_1567, which restored aminoglycoside efficacy against a diverse panel of multidrug-resistant Gram-negative ESKAPE pathogens.

### Identification of Potentiators of Aminoglycoside Activity

Our initial compound screen using *V. cholerae* reporter strain Naph3 led to the identification of eight compounds that synergise or potentiate kanamycin activity. Of these, three molecules (BMM_1567, BMS_4221, and BMM_2617) demonstrated robust potentiating effects at low concentrations with aminoglycosides across multiple resistance backgrounds, including the APH, ANT, and AAC resistance determinants. Notably, BMM_1567 showed potent activity at the lowest concentration range, suggesting both high efficacy and potential for lower toxicity. Its dual activity, potentiation at lower concentrations and synergy at higher doses, indicates a concentration-dependent mechanism of resistance inhibition that warrants further exploration.

### Activity Against Clinical MDR Isolates

When extended to a diverse panel of MDR clinical isolates from the ESKAPE group, BMM_1567 again emerged as the most promising candidate. It restored the activity of spectinomycin against MDR isolates of *E. coli*, *K. pneumoniae*, *A. baumannii*, and *P. aeruginosa*, achieving 50–100% growth inhibition at concentrations as low as 1.25 µM. Importantly, this compound also potentiated gentamicin and kanamycin activity against several resistant strains, although pathogen-specific variations in response were noted. For instance, BMM_1567 was effective with gentamicin against *E. coli* and *A. baumannii* but not *K. pneumoniae*, likely reflecting differences in permeability, efflux activity, or the specific resistance enzyme expressed.

BMS_4221 also showed activity, but required significantly higher concentrations and was less effective against *A. baumannii* and *P. aeruginosa*. Meanwhile, BMM_2617, despite its β-lactam structure, did not enhance activity against most β-lactamase-expressing isolates, limiting its utility as a potentiator in the current context (32, 33).

### Mechanism of Action: Targeting Resistance Enzymes

The selective potentiating activity of BMM_1567, in the absence of direct antibacterial effects, supports a mechanism involving inhibition of aminoglycoside resistance enzymes. Computational modelling provided strong evidence that BMM_1567 binds to the active sites of ANT, APH, and AAC enzymes, with the strongest affinity observed for ANT (ΔGbind ≈ - 62.25 kcal/mol). Importantly, interaction fingerprints showed significant overlap between BMM_1567 and spectinomycin binding sites within ANT, suggesting competitive inhibition with enzyme function. Similar trends were observed for APH and AAC, albeit with lower binding energies, indicating broader but variable efficacy across aminoglycoside-modifying enzymes. Similar trends have been observed for numerous β-lactamase inhibitors that have different efficacy against different β-lactamase classes (34).

To validate these interactions functionally, we employed site-directed mutagenesis targeting key ANT residues involved in BMM_1567 binding but not spectinomycin interaction (35, 36). The Leu168 and Trp169 substitutions significantly impaired resistance activity and diminished BMM_1567 potentiation. Notably, Trp169Tyr retained partial resistance while Leu168Ala resulted in a greater reduction in FICI, underscoring the importance of these residues in potentiator binding and enzyme inhibition. These findings not only indicate the potential mechanism of action of BMM_1567 potentiation but also provide a rational basis for future structure-guided optimization.

### Resistance Suppression and In Vivo Validation

The utility of any antibiotic adjuvant depends not only on its efficacy but also on its ability to suppress resistance emergence (37, 38). In our high-density plating assays (10⁸ CFU/mL), no resistant colonies were detected with the BMM_1567–spectinomycin combination. This result is particularly encouraging, suggesting that BMM_1567 may reduce the likelihood of resistance development by restoring antibiotic lethality at lower doses.

The efficacy of BMM_1567 was further confirmed in a murine skin infection model using an XDR *E. coli* isolate. Topical administration of BMM_1567 with spectinomycin significantly reduced bacterial burden by 2–3 log units compared to controls, rivalling the efficacy of colistin, a last-resort antibiotic. Importantly, no significant changes in body weight or pro-inflammatory cytokines (IL-6 and IL-1β) were observed, indicating safety and anti-inflammatory benefits of the combination therapy. Re-isolation and in vitro reassessment of bacteria from treated lesions confirmed that BMM_1567 retained its potentiating activity without selecting for resistance, further supporting its clinical potential.

### Broader Implications and Future Directions

This study highlights the promise of adjuvant-based therapeutic strategies to combat antibiotic resistance (39). BMM_1567, a Phase I/IIa clinical development peptide, demonstrates broad-spectrum potentiating activity against multiple aminoglycosides and MDR clinical isolates, while also exhibiting the clear molecular mechanism of potentiation action through inhibition of aminoglycoside-modifying enzymes. Very recently, BMM-1567 (LTX-315), has been reported to be associated with membrane-targeting antibacterial activity at elevated concentrations (40). In the present study, we demonstrate that the compound functions as an aminoglycoside potentiator by inhibiting the resistance enzyme. The efficacy of BMM-1567 as aminoglycoside potentiator in a murine model underscores its translational relevance.

Several key future directions arise from this work. First, pharmacokinetic and toxicity profiling of BMM_1567 will be essential to determine suitability for systemic use. Second, structural analogs of BMM_1567 can be designed based on our interaction data to further enhance potency and spectrum. Third, combination strategies involving BMM_1567 with other antibiotic classes or efflux pump inhibitors may yield synergistic effects in particularly recalcitrant pathogens like *P. aeruginosa*. Finally, the observation that different compounds display pathogen-specific and enzyme-specific activities highlights the importance of personalized adjuvant therapies, possibly guided by rapid diagnostics to identify resistance genes.

## Conclusion

In summary, our study establishes aminoglycoside-modifying enzymes as therapeutically actionable targets for overcoming AMR and identifies BMM_1567 as a first-in-class aminoglycoside potentiator capable of restoring antibiotic efficacy against diverse MDR Gram-negative pathogens. By selectively disabling resistance mechanisms rather than directly targeting bacterial viability, BMM_1567 enhances antibiotic activity, suppresses the emergence of resistance, and improves therapeutic outcomes in vivo. More broadly, this work provides a scalable framework for the discovery of resistance-targeting antibiotic adjuvants and demonstrates how integrating genome-guided phenotypic screening, mechanistic validation, and computational modeling can uncover new therapeutic strategies against AMR. Given the urgent need for approaches that extend the lifespan of existing antibiotics, resistance-enzyme inhibition represents a promising paradigm for revitalizing the clinical utility of legacy antimicrobials and combating the global threat of multidrug-resistant bacterial infections.

## Methods

### Bacterial strains and growth conditions

The *V. cholerae* strain N16961, and *E. coli* ATCC25922 strains were used in this study. We have constructed multiple reporter strains using N16961, and *E. coli* ATCC25922 strains. All the strains constructed and used are mentioned in **Table 1**. All the genes cloned, their source and plasmid constructed are mentioned in **Table S2.** For liquid culture, the strains were grown in Luria Broth (LB), Muller-Hinton Broth (MHB) medium at 37°C in a shaker with 180 rpm while LB agar plates and Muller-Hinton agar (MHA, Difco, USA) were used for solid culture. The following antibiotics were used: streptomycin, spectinomycin, kanamycin, and zeocin. *V. cholerae* strains were routinely stored on solid agar plates at room temperature. *E. coli* strains were routinely stored on solid agar plates at 4°C. For long term storage at –80°C, 15% glycerol supplemented LB medium was used.

### Molecular biological methods

Unless otherwise specified, conventional molecular biology procedures were used for chromosomal and plasmid DNA isolations, electroelution, restriction enzyme digestion, ligation of DNA fragments, bacterial transformation, conjugation, and so on (22, 41). All restriction enzymes and nucleic acid-modifying enzymes were sourced from New England BioLabs, Inc. and applied in accordance with manufacturer’s instructions. Transformants were chosen by plating transformed cells on LB agar plates with antibiotics.

### Development of antibiotic resistance plasmids and reporter strains

Construction of resistance plasmids containing antibiotic resistance genes were carried out by using the integrative conjugative vector pBD62 and its derivative pSB49 (23). The MDR and XDR clinical isolates were used as templates to amplify ARGs (**Table S2)**. Using gene specific primer combinations carrying restriction enzymes, the corresponding resistance genes either with its native promoter or the ORF alone were PCR-amplified (**Table S2)**. The amplified components were purified even before restriction enzyme digestion, and ligated into likewise digested vectors pBD62 and pSB49. For expression under inducible promoter (P_BAD_) the in-frame cloning was done in pBD62 and for expression under constitutive promoter (P_htpG_) the in-frame cloning was done in pSB49. The host bacteria *E. coli* FCV14 was utilized to select and replicate recombinant vectors. The recombinant vectors were transferred to *V. cholerae,* and *E. coli* ATCC via conjugation. The donor strain used was *E. coli* β2163. The reporter strains were selected on respective antibiotics and further confirmed by PCR.

### Minimum Inhibitory Concentration (MIC) determination

MIC of reporter strains were determined by broth dilution method of antimicrobial susceptibility testing according to CLSI guidelines (42, 43). In brief, overnight bacteria cultured in MHB medium were diluted at 1:100 in fresh MHB and cultured at 37°C with shaking at 180 rpm to an optical density of 0.5(2 × 10^8^ CFU/ml) measured at 600nm. Then, diluted at 1:1000 in fresh MHB (2 × 10^5^ CFU/ml). Then 100μL of this diluted culture was added into each well of a 96-well microtiter polystyrene tray. A series of 2-fold dilutions of an antibiotic was made in this 96-well plate. The mixtures were incubated at 37°C for 16-18h. MIC was defined as the lowest antibiotic concentration that inhibited visible bacteria growth.

### Screening of Compound libraries

#### Primary Screening

Overnight reporter strains cultured in MHB medium were diluted at 1:100 in fresh MHB and cultured at 37°C with shaking at 180 rpm to an optical density of 0.5 (2 × 10^8^ CFU/ml). Then, diluted at 1:1000 in fresh MHB to achieve an initial inoculum of 2 × 10^5^ CFU/ml. The 145μL of this diluted culture with test antibiotic was added into each well of a 96-well microtiter polystyrene tray. 5μL from 300μM stock solutions of each compound dissolved in appropriate solventswas added to each separate well of a 96-well plate. The controls of screening include bacterial culture alone, bacterial culture with test antibiotic, bacterial culture with both test antibiotic and the solvent, bacterial culture with sensitive antibiotic and blank media control. The screening was done in triplicates. The mixtures were incubated at 37°C and after 16-18 hours the optical density at 600 nm (OD600) of the plate was determined using a spectrophotometer. The % inhibition was calculated as (PC^Ab^-Test^Ab^/PC^Ab^-Neg)*100 where PC is bacterial culture with the antibiotic and DMSO, Test is bacterial culture with antibiotic and the compound and Neg is the MHB alone.

#### Secondary screening

The compounds from primary screening were further assessed through secondary screening. Following the same protocol for bacterial inoculum preparation as previously described, a 96-well plate was prepared. Each well contained bacterial culture with test antibiotics, and an additional set contained only bacterial culture. To these wells, 5μL from 300μM stock solutions of each compound was added. The mixtures were incubated at 37°C and after 16-18 hours, OD600 of the plate was determined using a spectrophotometer. Potentiating or synergistic compounds that displayed minimal or no antimicrobial activity and inhibited at least 30% bacteria growth in combination with test antibiotics, were identified for further consideration. The secondary screening was done with and without antibiotic and % inhibition was calculated.

#### Checkerboard assay

Overnight reporter strains (E-aph3’, E-aadA1, E-aac3) cultured in MHB medium were diluted at 1:100 in fresh MHB and cultured at 37°C with shaking at 180 rpm to an optical density of 0.5 (2 × 10^8^ CFU/ml). Then, diluted at 1:1000 in fresh MHB to achieve an initial inoculum of 2 × 10^5^ CFU/ml. The 100μL of this diluted culture was added into each well of a 96-well microtiter polystyrene tray. In the first row and last column 200μL was added. Add antibiotic and compound at concentration double the MIC. The test antibiotic in the combination is diluted sequentially along the ordinate, while the compound is diluted along the abscissa. The resulting checkerboard contains each combination of antibiotic and compound. The plate was incubated at 37°C and after 16-18 hours, OD600 of the plate was determined using a spectrophotometer.

In order to quantify the interaction between the tested antibiotics (FIC index), the following formula was used:

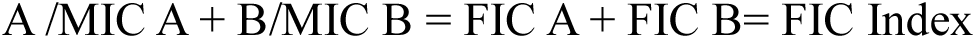

Where A and B are the MICs of antibiotic and compound combined, and MIC A and MIC B are the MICs of antibiotic and compound, respectively. The FIC Index value is calculated to classify the interaction of the antibiotics and compounds. When the FIC value is less than 0.5, the effect is synergistic; when FIC> 4, the effect is antagonistic; when the FIC is 0.5-4, the two are represented as indifference or additive.

#### Molecular docking

The sequences of Aminoglycoside-modifying enzyme (ANT), Aminoglycoside phosphotransferase (APH) and Aminoglycoside acetyltransferase (AAC) were retrieved from uniprot. In absence of crystal structures of these protein the template-based homology modeling was carried out using pdb-ids: 7UY4, 4FEU and 7LAO for ANT, APH and AAC, respectively. The crystal structures were retrieved from the RCSB Protein Data Bank (www.rcsb.org). The built models were prepared using the Protein Preparation Wizard module of Maestro (Schrödinger Release 2022-1: Maestro, Schrödinger, LLC, New York, NY, 2022). In the preparation process, the hydrogen and bond orders were added using PRIME. The hydrogen bond (HB) optimization and restrained minimization was also done for the systems using the OPLS3 force field model. Since the template co-crystal structures (bound with antibiotics) are close homologs of ANT, APH, and AAC, hence these crystals were used for structural and binding comparison. The molecular docking of BMM_1567 was carried out using the HDOCK. The structure of peptide was prepared using the Protein Preparation Wizard module of Maestro (Schrödinger Release 2022-1: Maestro, Schrödinger, LLC, New York, NY, 2022) (44).

#### Molecular dynamics (MD) simulations

After docking the top complexes were analyzed and the most likely pose was quantified using MM-GBSA and molecular dynamics (MD) simulation studies (45). The binding free energy of all three protein-peptide complexes was carried out in prime MM-GBSA (molecular Mechanics Generalized Born Surface Area). The binding free energies of the protein-peptide complex were calculated using the Prime MM-GBSA module of Schrodinger Suite with an OPLS4 force field. The MD simulation was carried out in DESMOND module of Schrodinger suite using OPLS4 force field. Total 1.8μs MD simulations of different systems (3 protein-peptide, 3 protein-antibiotic, and 3 APO) were carried out to compare the dynamics difference between APO, Pro-pep and Pro-drug systems using post-processing analysis of MD trajectories. First, the simulation systems were build using the OPLS4 force field and the protein complexes using the TIP3P water model. An orthorhombic box shape was chosen, with an edge of 12.0 Å ensuring the minimal distance between the atoms of protein and the edge of the box. To neutralize the system, counter ions were added. An energy minimization of each system was done using steepest descent integrator for 2000 steps. For the simulation, the NPT ensemble was used at 300k in temperature and 1.01325 bar in pressure. The simulation was conducted for 200 ns for each system, and trajectories were saved every 20ps. Desmond analysis modules were used for post-processing analysis. The average binding free energies were calculated for MD trajectories for protein-peptide complexes of all three proteins using MM-GBSA approach. For the calculations, the stable frame was extracted from the MD trajectory.

### Animal studies

#### Ethics statement

All animal experiments were performed after obtaining ethical approval from the Institutional Animal Ethics Committee of Translational Health Science and Technology Institute (THSTI) with approval no. – IAEC/THSTI/352.

#### Skin abscess infection mouse model

The potentiating activity of the lead compound with antibiotic was assessed in vivo using a skin abscess mouse model against an XDR *E. coli* strain, following reported methodology (46). To assess the effectiveness of the potentiator-antibiotic combination against XDR *E. coli* strain, the bacteria were cultured in MHB medium until reaching an OD_600_ of 0.3. Subsequently, the cells were washed twice with sterile PBS (pH 7.4) and suspended to make a concentration of 7×10^6^ colony-forming units CFU/ml. For the in vivo experiments, female BALB/c mice (7 to 8 weeks old) were grouped as follows - untreated, vehicle (1.5% DMSO), spectinomycin alone (80µg/ml), BMM_1567 alone (2.5µM), spectinomycin + BMM_1567 (80µg/ml+2.5µM), BMS_4221 alone (50µM), spectinomycin + BMS_4221 (80µg/ml+50µM) and colistin treated (20µg/ml). The hair of the mice was trimmed and shaved at the area of interest and their skin was disinfected with povidone iodine solution. Using sterile dissecting scissors and forceps, the top layer of skin on the back was then removed, creating an incision of around 1 cm. The wound was inoculated with a 20µL aliquot of the bacterial solution suspended in PBS. One hour after infection, the initial dose was applied to the affected area for all groups, and followed by a second dose on day 1. Two days post-infection, the animals were euthanized, and the skin area with the infection was excised and homogenized using a bead beater. The homogenates were then 10-fold serially diluted and plated on spectinomycin (40µg/ml) plate for CFU quantification. All the groups had 5 mice per group except for vehicle group which had 3 mice.

#### Real time reverse transcriptase PCR for expression of IL-6 and IL-1β

Blood was collected from the mice via retro-orbital bleeding after two days post-infection and total RNA was extracted using the Qiagen Mini Kit following the manufacturer’s protocol. First strand cDNA was synthesized using Qiagen’s Quantitect Reverse Transcription kit from 900 ng RNA. Real time PCR was carried out in a QuantiStudio^TM^ 6 Real time PCR machine of Thermo Fisher Scientific using the PowerTrack SYBR Green Master Mix (AppliedBiosystem). The primers used are listed in **Table S3**. Glucose-3 phosphatedehydrogenase (GAPDH was used as the indigenous control.)

#### Prevalence of ARGs

Genomes of *A. baumannii*, *E. coli, K. pneumoniae* and *P. aeruginosa* available on NCBI were downloaded with the following filters: host = *Homo sapiens,* isolation source = clinical, collection year = 2019-2023 for *A. baumannii*, *E. coli, K. pneumoniae* and 2014-2023 for *P. aeruginosa*. The CARD database in the ABRicate tool was used to predict the ARGs in the isolates with a filtering criterion of 80% identity and 80% gene coverage.

##### Data availability

All data generated and analyzed during this study are original and are included in this article and its Supplementary Information files. Additional data supporting the findings of this study are available from the corresponding author upon reasonable request.

## Author contributions

BD conceived the idea and designed the experiments. MC, LN, DP, RSK, SK, DD, KK, SB, PP conducted the experiments. MC, LN, SK, DP, DM, SA, BD, performed data analysis. MC, LN, DP, SA and BD wrote the manuscript. BD edited the manuscript. All authors have read and approved the manuscript.

### Disclosure and competing interests statemen

The authors declare no competing interests.

## Acknowledgements

Ms. Meenal Chawla is thankful to CSIR, Govt. of India for the research fellowships. Dr. Lekshmi N and Dr. Deepjyoti Paul acknowledge the support of the Department of Biotechnology, Govt. of India for the MK Bhan fellowship program for research support. The authors thank all the technical staff of the Advanced Nucleotide Sequencing Facility at THSTI for their technical support in sequencing the clinical isolate genomes. The authors would like to express their sincere gratitude to Prof. Ganesan Karthikeyan, Executive Director of the BRIC-Translational Health Science and Technology Institute (THSTI), for his unwavering support in helping to complete this study.

